# The variability of reflex amplitude estimates in motor unit pools depends on the phenotype distribution and discharge statistics

**DOI:** 10.64898/2026.02.11.705250

**Authors:** Laura Schmid, Thomas Klotz, Oliver Röhrle, Christopher K. Thompson, Francesco Negro, Utku Ş. Yavuz

**Affiliations:** Institute for Modelling and Simulation of Biomechanical Systems, University of Stuttgart, Stuttgart, Germany; Stuttgart Center for Simulation Sciences (SC SimTech), University of Stuttgart, Stuttgart, Germany; Department of Health and Rehabilitation Sciences Temple University, Philadelphia, Pennsylvania, USA; Department of Clinical and Experimental Sciences, Università degli Studi di Brescia, Brescia, Italy; Department of Biomedical Signals and Systems, Faculty of Electrical Engineering, Mathematics and Computer Sciences, University of Twente, Enschede, Netherlands

**Keywords:** peristimulus time histogram, peristimulus frequencygram, H-reflex, electromyography, motor unit decomposition, mathematical model, computer simulation

## Abstract

Motor unit (MU) activity during electrically or mechanically evoked reflexes is used to examine the relationship between neural inputs and MU properties. However, variations in single-MU reflex amplitudes are not fully understood and limit their reliability in determining the input-output relation of motor neurons (MNs). Using experiments and computer simulations, we investigated (i) whether MN discharge statistics and muscle activation explain the variability of reflex amplitude estimates and (ii) whether these variations are reflected differently across distinct reflex amplitude estimation methods.

We analyzed MU spike trains extracted from isometric contractions of the tibialis anterior muscle at 10 % and 20 % MVC (maximum voluntary contraction). Estimating reflex amplitudes based on the peristimulus frequencygram (PSF) at 10 % MVC, the linear regression between discharge rate (DR) and reflex amplitude was always positive, with *p* < 0.05 in 3 out of 6 subjects; however, the linear correlation was inconsistent at 20 % MVC. We thereby observed that inter-subject variability was associated with the coefficient of variation of the interspike intervals. Furthermore, the linear correlation between DR and peristimulus time histogram (PSTH) based reflex amplitudes was inconsistent for both contraction forces.

To obtain further insights into the influence of MN properties, we simulated reflexes in a heterogeneous MN population using electrical circuit models and varied MN inputs. The simulations indicate that, besides mean input current and membrane noise, MN properties also contribute to the variability of reflex amplitude estimates. The MN heterogeneity is well captured by PSF-based reflex estimates but not by PSTH-based ones. These results show that variations in amplitude estimates of individual MU reflexes are due to complex interactions between intrinsic and extrinsic factors. As PSF-based reflex amplitude estimates reflect the MN size distribution, tracking PSF-based reflex amplitudes at fixed MVC levels across individual subjects could serve as a marker for investigating spinal adaptations under (patho)physiological conditions.

**Author summary:** Motor neurons are specialized nerve cells that control human movement. Each motor neuron activates a specific set of muscle fibers, and the functional unit consisting of a motor neuron and muscle fibers is called a motor unit. We can observe the activity of motor neurons in humans by decomposing the electrical activity of muscles (the electromyogram) into contributions from individual motor units. Reflex responses of motor units are often used to study the input-output relation of motor neurons in humans. We used a combination of experiments and computer simulations to study the factors that influence the reflex amplitude of motor units during an excitatory reflex. We found that the reflex amplitude is non-linearly influenced by a number of intrinsic and extrinsic factors, e. g., motor neuron size, but also the muscle force. Additionally, we found that these factors have different effects on the results of the two common methods used to calculate the reflex amplitude. These results provide guidance on choosing a suitable evaluation method and on interpreting reflex experiments.

## 1 Introduction

Due to its simple monosynaptic projection, the mechanically or electrically evoked Ia reflex is commonly used to study the input-output relation of spinal (alpha) motor neurons (MNs) in humans. The reflex amplitude of a single motor unit (MU) is believed to be related to the postsynaptic potential (PSP), and the gain of this relation is assumed to represent MN excitability (Powers and Türker, 2010). Based on this assumption, the reflex amplitude of MUs can be used to estimate the excitability of spinal MNs and their adaptation to specific physiological or pathological conditions (Scaglioni et al., 2003; Norton et al., 2008). However, excitability is not the only factor that affects the reflex amplitude. Several extrinsic factors, including recurrent and presynaptic inhibition, as well as excitatory and inhibitory postsynaptic inputs, contribute to fluctuations in the reflex amplitude (Özyurt et al., 2019; Morita et al., 1998). In addition to extrinsic factors, several intrinsic factors, including the firing regime of a MN itself, were hypothesised to contribute to the reflex amplitude. For example, the background DR and random membrane potential fluctuations affect MU reflex amplitude estimates (Jones and Bawa, 1995; Powers and Türker, 2010; Poliakov et al., 1996). These observations are also supported by computer simulations demonstrating that the response probability of an individual MN to an excitatory or inhibitory postsynaptic potential (EPSP or IPSP) varies with the mean DR and membrane noise (Jones and Bawa, 1997; Herrmann and Gerstner, 2002; Schmid et al., 2024). Moreover, due to the stochastic behaviour of neurons, single MN reflex estimates are associated with estimation errors that depend on the number of applied stimuli (Schmid, 2024).

The discharge behaviour of a MU in response to a reflex is often investigated using two distinctive methods: the peristimulus time histogram (PSTH) and the peristimulus frequencygram (PSF). The PSTH is based on counting spikes in time bins (Ellaway, 1978), whereas the PSF computes the instantaneous firing frequencies of the spikes (Türker and Cheng, 1994). Both methods require delivering multiple reflex stimuli to statistically analyse MU activity relative to the timing of the reflex stimulus. Due to the methodological differences, the estimated reflex trajectories may differ for the same MU (Türker and Powers, 2005). Many studies have addressed the synchronization-related erroneous interpretation of the PSTH, which can be avoided by using the PSF (e. g., Türker and Powers, 2005; Yavuz et al., 2014). Türker and Powers (1999) showed in their *in-vitro* study that PSTH and PSF reflect different aspects of the PSP, e. g., the shape or size.

Studies investigating single motor neuron firing patterns in humans typically exploit the selective properties of the intramuscular electromyogram (iEMG) using template-based signal decompositions (McGill et al., 2005). With the development of sophisticated blind source separation methods (Negro et al., 2016a; Holobar and Zazula, 2007), high-density surface EMG recordings can be decomposed into tens of MU spike trains during reflex experiments (Yavuz et al., 2015). A larger sample of MUs allows us to investigate the influence of DR and membrane noise on H-reflex responses with a stronger statistical outcome and provides the possibility of using population analysis to unveil so far unknown patterns. We hypothesize that understanding the multivariable variation in the reflex amplitude can enable us to better estimate the adaptation in MN pools in physiological and pathological conditions.

In the present study, we combine experimental measurements and computer simulations of MN pools to investigate whether the DR statistics of a sampled MN pool can explain the variation in reflex amplitude distribution and to what extent the chosen analysis method influences our interpretations. The simulation results augment the database available for the analysis and enable unique insights that would not be easily feasible experimentally. We hypothesized that the distribution of the H-reflex amplitude of a MU population will be influenced by the MU discharge properties, i. e., DR, coefficient of variation of the interspike interval (CoV_ISI_) and the MN size. The variation in the reflex amplitude due to discharge regime and other factors needs to be taken into account before interpreting reflex amplitude as a measure of excitability.

## 2 Results

### 2.1 Influence of discharge statistics

#### *In-vivo* results

The mean stimulation intensity to evoke the H-reflex was 10.9±3.2 mA. The total number of identified MUs was 206 in two modalities (176 from HDsEMG and 30 from iEMG) and at two contraction force levels (10 % and 20 % MVC). A summary of the reflex analysis (PSF-based and PSTH-based reflex amplitude, latency and duration) and MU firing statistics (background DR and firing rate variability) for the two MVC levels is given in Table 1. As expected, the reflex latency (approx. 40 ms) and duration (approx. 10 ms) were not different between contraction forces (*p* > 0.05). Further, the mean DR was higher at 20 % MVC (*p* < 0.001, *F* = 20.022), while the CoV_ISI_ was not significantly different between contraction forces (*p* > 0.05). Increasing the contraction intensity decreases the reflex amplitudes, i.e., comparing the mean values at 20 % and 10 % MVC, we observe a decrease by 27 % for PSF (*p* = 0.040, *F* = 8.551) and 21 % for PSTH (*p* = 0.048, *F* = 3.96).

**Table 1:** The reflex parameters and MU discharge statistics at 10 % and 20 % MVC obtained from the experimental study. Asterix (*) indicates significant differences.

| Property | Contraction level |  |
| --- | --- | --- |
|  | 10 % MVC | 20 % MVC |
| Number of MUs | 106 | 100 |
| Latency (ms) | $41.51 \pm 4.93$ | $41.41 \pm 5.22$ |
| Duration (ms) | $10.64 \pm 6.77$ | $9.55 \pm 6.63$ |
| Reflex amplitude PSF (Hz/N.Stim.) | $1.45 \pm 1.13^*$ | $1.06 \pm 0.74$ |
| Reflex amplitude PSTH (count/N.Stim) | $0.24 \pm 0.18^*$ | $0.19 \pm 0.16$ |
| mean DR (Hz) | $10.31 \pm 1.31$ | $11.17 \pm 1.42^*$ |
| CoV<sub>ISI</sub> (%) | $14.5 \pm 5.4$ | $15.0 \pm 6.0$ |

We fitted a linear regression model (for each subject at each MVC level) to test whether the variability in DR can explain the variability of the measured reflex amplitudes. Figure 1 shows the regression models for six subjects during H-reflex at 10 % and 20 % MVC. The shaded areas along the regression models depict unexplained variations in reflex amplitude, so-called residuals. At 10 % MVC, for all subjects, we found a positive linear relation between PSF-based H-reflex amplitude and mean DR, which reached the significance threshold (*p* < 0.05) in 3 out of 6 subjects. In contrast, at 20 % MVC, 3 out of 6 subjects showed a negative linear relation and 3 out of 6 subjects showed a positive linear relation between DR and PSF-based reflex amplitude. However, for all subjects and 20 % MVC, the linear relations are not significant (*p* > 0.05). We found a weak negative linear correlation between PSTH H-reflex amplitude and mean-DR (4 out of 6 subjects at 10 % MVC and 3 out of 6 subjects at 20 % MVC); however, the correlations are not significant (*p* > 0.05).

**Figure 1.**
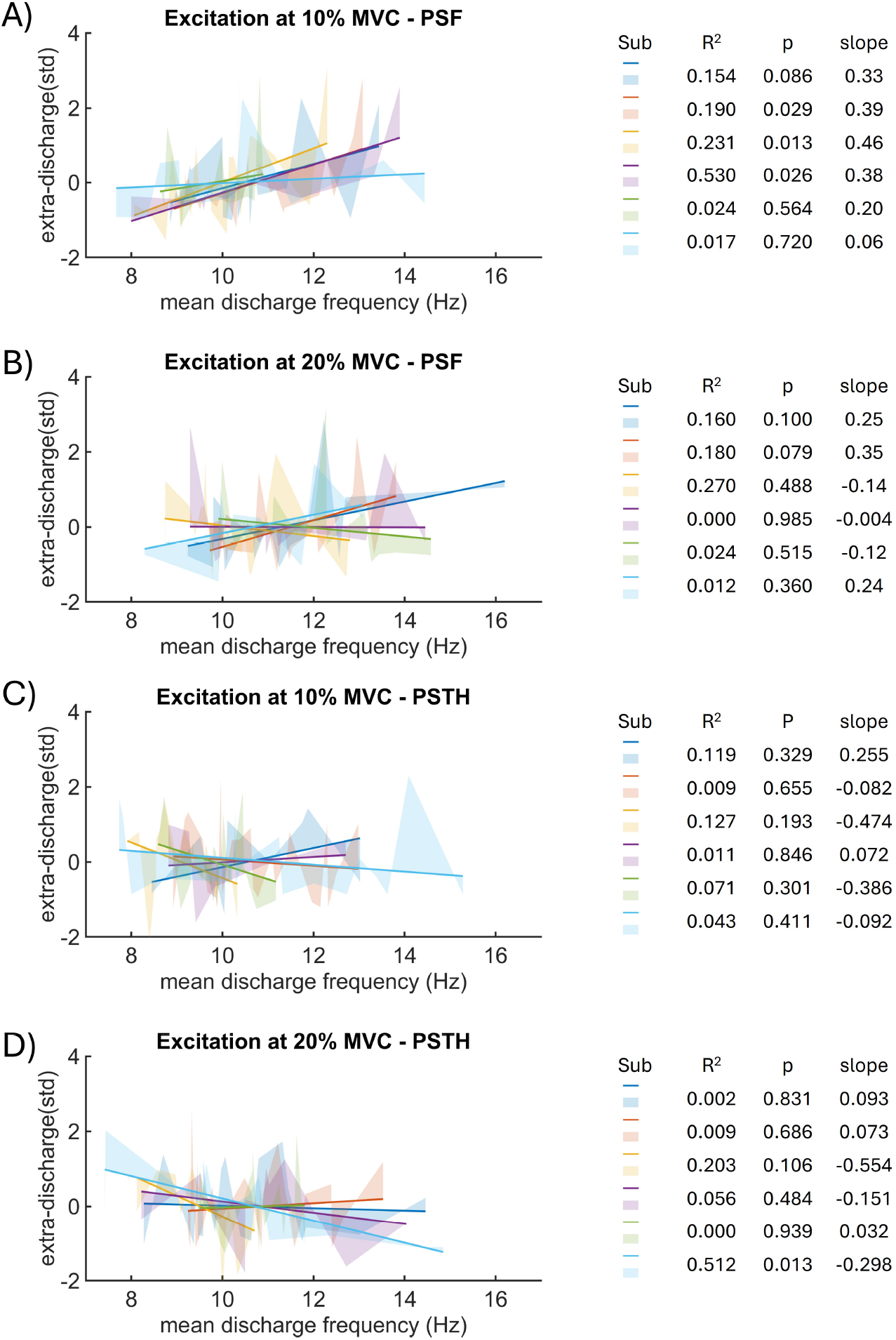
Linear regression between background DR and reflex amplitude for six subjects. Solid lines show the linear regression model, and shaded areas show residuals. For each subject, the coefficient of determination (R^2^), the p-value (p), and the slope of the regression model are given. Shown are the reflex amplitudes determined from PSF at 10 % (A) and 20 % (B) MVC, as well as reflex amplitudes determined from PSTH at 10 % (C) and 20 % (D) MVC.

We examined whether the inter-subject variability of the PSF-based regression models is associated with membrane noise, which is quantified by the variability between spike intervals (CoV_ISI_). To this end, subjects were divided into subgroups based on mean discharge rate (DR) and reflex amplitude regression results estimated at 10 and 20 % MVC. Group 1 involved subjects who showed significant or consistently positive regression between DR and reflex amplitude, while Group 2 consisted of the remaining ones. We found a significant positive linear relationship between CoV_ISI_ and reflex amplitude (*p* << 0.001) for the group that also has a significant positive correlation between DR and reflex amplitude at 10 % MVC (Figure 2). This group also showed lower CoV_ISI_ values compared to the group that has no significant linear relation between DR and reflex amplitude. This result suggests that the higher CoV_ISI_ and thus membrane noise disturb the correlation between DR and reflex amplitude. These results need to be interpreted under the assumption of the independence of mean DR and ISI variability. Our results show a weak correlation between mean DR and CoV_ISI_ at both 10 % (r=0.35) and 20 % MVCs (r=0.27).

**Figure 2.**
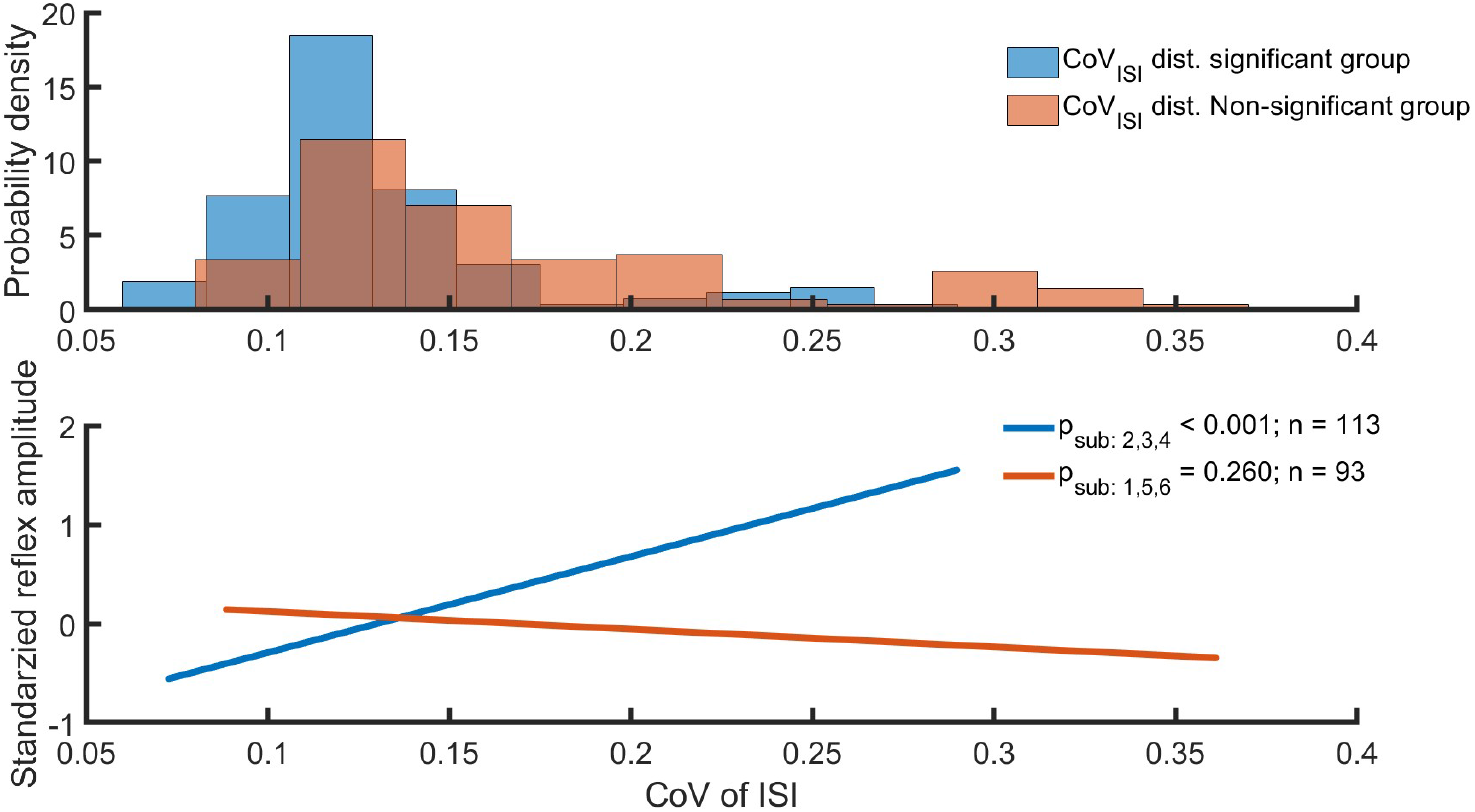
The relation between the reflex amplitude and the coefficient of variation of inter-spike-interval (CoV_ISI_) based on inter-subject variability. Subjects were divided into subgroups based on mean discharge rate (DR) and reflex amplitude regression results. Blue label shows results from group 1 (subjects 2, 3, and 4), which showed significant or consistently positive regression. Orange is the result of the group 2 (subjects 1, 5, and 6) without a significant DR-Reflex amplitude regression. A: The probability density distribution of the CoV_ISI_. B: Linear regression between reflex amplitude and CoV_ISI_.

#### Simulation results

Similar to the *in-vivo* MU data, we fitted a linear regression model between the DR and the standardized reflex amplitudes for the simulated MNs. In the simulated MN population, DR ranges from 7.05 Hz to 38.83 Hz. Note that in the simulation, we only considered one MN pool consisting of 200 MNs and used seven (instead of two) activation levels, corresponding to physiological force levels from approx. 5% MVC to full recruitment. Figure 3A shows the number of recruited MNs at the chosen activation levels as well as their input resistance. In the simulation, 44.5 % of MNs are recruited at the lowest activation level 1 and 98.5 % at the highest activation level 7. The current-frequency relations of the 200 simulated MNs are shown in Figure 3B. With increasing activation (representing effective synaptic input), more MNs are successively and orderly recruited according to the size principle (Henneman et al., 1965a,b). All MNs show increasing firing rates for increasing effective synaptic input. The smallest MN (MN 1) increases its mean DR from 10.12 Hz to 38.72 Hz and MN 197 is the largest recruited with 11.06 Hz at activation level 7. The slope of the current-frequency relation is not constant, but the initial increase is steeper than the later one. Table 2 shows the number of MNs with significant reflexes using the PSF-based and PSTH-based methods, respectively. The reflex responses estimated from the PSTH were significant for almost all MNs (except activation level 4). The relative number of significant reflexes measured in the PSF decreased with increasing activation. Note that MNs with non-significant reflex responses were excluded from the following analysis.

**Figure 3.**
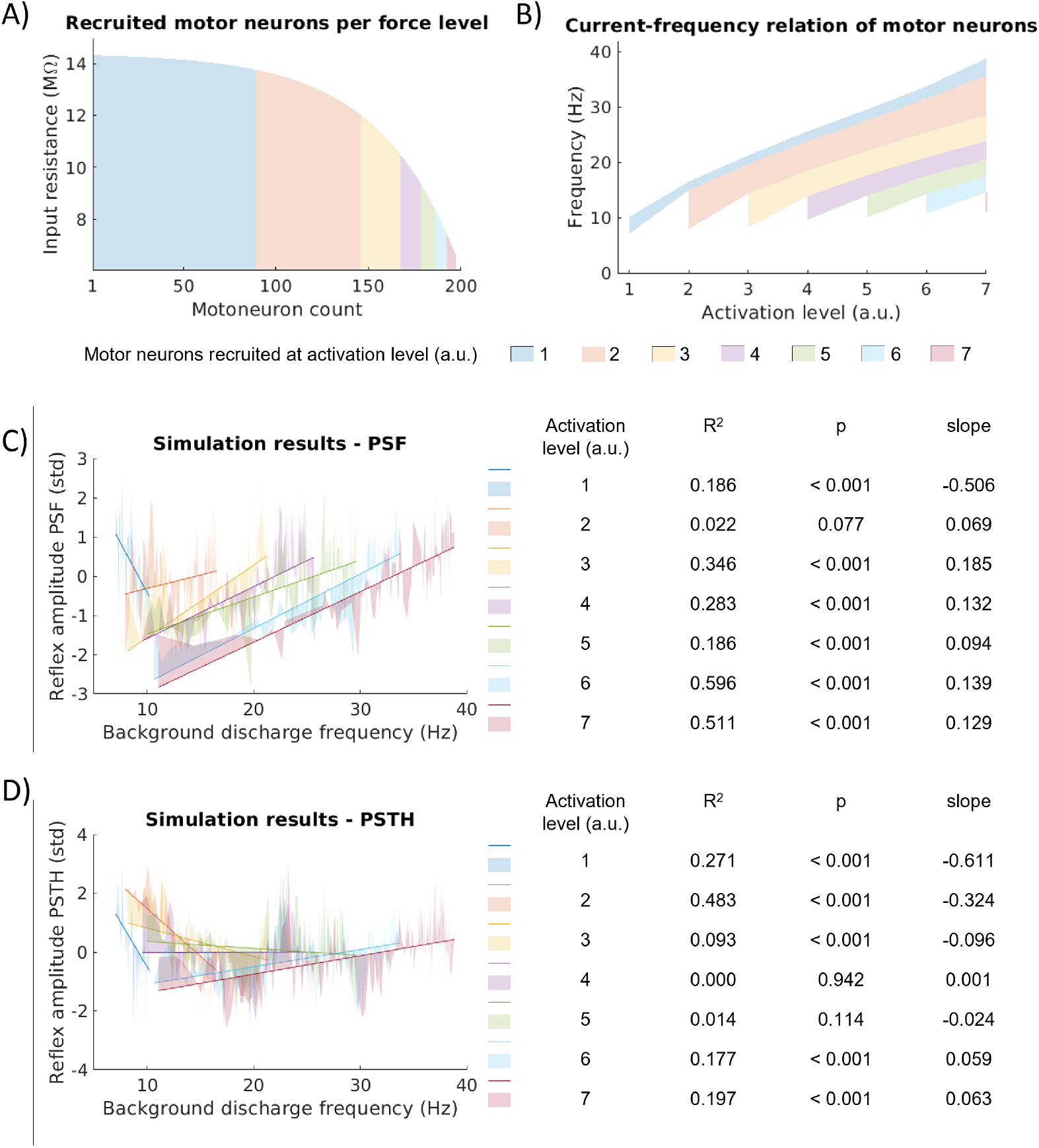
A: Input resistance and number of additionally recruited motor neurons per activation level for the employed simulation setting. B: Current-frequency relation of the respective groups of motor neurons. C: Standardized reflex amplitude determined from PSF. d: Standardized reflex amplitude determined from PSTH. For each activation level, the coefficient of determination (R^2^), the p-value (p) and the slope of the regression model are given.

**Table 2:** Number of included motoneurons (MNs) and number of MNs with a significant reflex amplitude depending on the activation level in arbitrary units (a.u.). Reflex amplitudes were determined from the cumulative sum of the peristimulus time histogram (PSTH) and the peristimulus frequencygram (PSF). Simulated MNs with background DR above 7 Hz and a CoV_ISI_ below 30 % were included.

| Property | Activation Level (a.u.) |  |  |  |  |  |  |
| --- | --- | --- | --- | --- | --- | --- | --- |
|  | 1 | 2 | 3 | 4 | 5 | 6 | 7 |
| Included MNs | 89 | 148 | 169 | 181 | 188 | 193 | 199 |
| Significant reflex in PSTH | 89 | 148 | 168 | 179 | 188 | 193 | 199 |
| Significant reflex in PSF | 89 | 148 | 168 | 174 | 151 | 146 | 91 |

The linear relation between PSF reflex amplitude and background DR is negative for activation level 1 and positive for the other activation levels (Figure 3c). The R^2^-values do not exceed 0.6. The linear regression model reaches significance (*p* < 0.05) for all but activation level 2. For the linear correlation between PSTH reflex amplitude and background DR, four of the seven activation levels are negative. The R^2^-values do not exceed 0.5 and the linear regression model reaches significance for all but activation levels 4 and 5.

### 2.2 Influence of MN size properties (simulation)

#### Tracking single motor neurons across activation levels

To investigate the relation between MN reflex amplitude and applied input in more detail, we tracked simulated MNs across seven activation levels (Figure 4A, B). The reflex amplitudes determined from the PSTH-cusum range from 0.073 counts/No. of Stim to 0.565 counts/No. of Stim and the reflex amplitudes determined from the PSF-cusum range from 0.108 Hz/No. of Stim to 2.017 Hz/No. of Stim. The relation of reflex amplitude and activation level differs between PSTH and PSF. In the PSTH, reflex amplitudes generally decrease with increasing activation and, finally, plateau. Thereby, the CoV of the reflex amplitude increases from 7 % to 23 %. With increasing activation, the PSF reflex amplitudes of small MNs first decrease and then increase. In contrast, the reflex amplitudes of larger MNs consistently decrease with increasing activation. The pool of MNs shows relatively similar reflex amplitudes for low activation levels (CoV of reflex amplitudes 10 % at activation level 1), but the range of values increases with increasing activation (CoV of reflex amplitudes 41 % at activation levels 6 and 7). While the totality of values shows a clear trend, individual MNs (dots connected by lines) generally do not show consistent behaviour, indicated by crossing lines.

**Figure 4.**
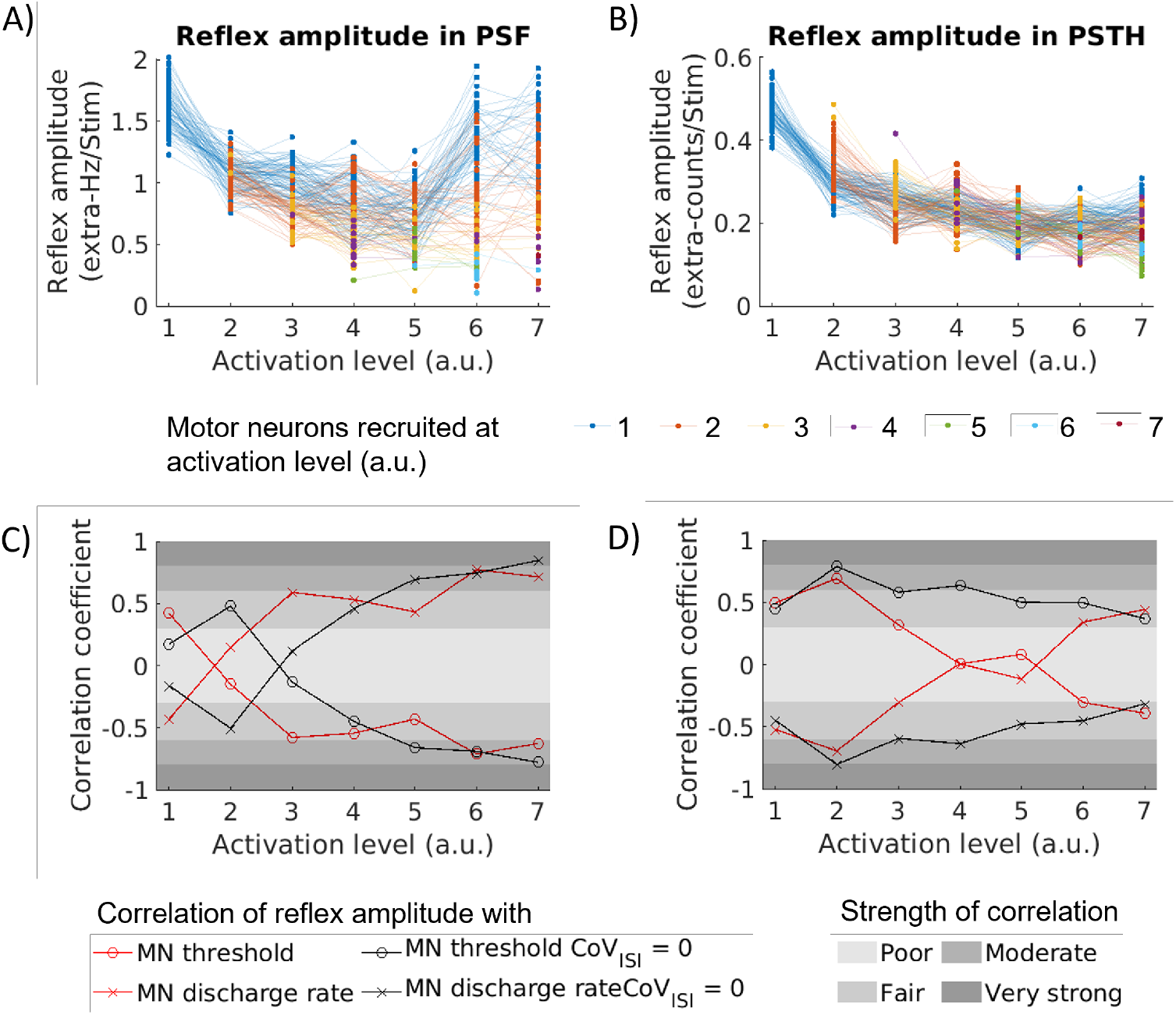
Reflex amplitude from PSF (A) and PSTH (B) for the pool of MNs and for each activation level. Reflex amplitudes of individual MNs are dots connected by lines. Colors correspond to Figure 3. Linear correlation coefficient of reflex amplitude from PSF (C)/PSTH (D) and MN recruitment threshold (o)/discharge rate (x). Red lines show the result for data shown in (A) and (B); the black lines correspond to simulations performed with CoV_ISI_ = 0. Rating of correlations according to Chan YH scale (Chan, 2003).

#### Linear correlation coefficient per activation level

We calculated the linear correlation coefficient (Pearson’s) between reflex amplitude and background DR as well as between reflex amplitude and recruitment threshold for both PSF and PSTH (Figure 4C, D). The PSF reflex amplitude correlates negatively with the DR for activation level 1 but positively and successively stronger for higher activation levels. The highest correlation coefficient is reached at activation level 6 (|*ρ*| =0.772, p <0.05), which can be considered a moderate correlation according to Chan YH scale (Chan, 2003). Except for activation level 2, which shows poor correlation (|*ρ*| =0.147, p = 0.077), the correlation is at least moderate for all other activation levels (|*ρ*| ≥ 0.432, p < 0.05). If the membrane noise is omitted (i. e., CoV_ISI_ = 0), the results look qualitatively similar but are shifted towards higher activation levels.

For the PSTH the trend is less consistent across activation levels. The correlation with DR is fair to moderate and negative for activation levels 1 to 3 (0.305 ≤ |*ρ*| ≤ 0.695, p < 0.05), then drops to values close to zero for activation levels 4 and 5 (|*ρ*| ≤ 0.116, p ≥ 0.114) and increases to positive values for activation levels 6 and 7 (|*ρ*| ≥ 0.343, p < 0.05). Overall, the correlations are weaker than in the PSF results. Omitting the membrane noise (i. e., CoV_ISI_ = 0) increases the correlation coefficients for all activation levels but 1 and 7. Correlations are fair to moderate for all activation levels, reaching a strong correlation for activation level 2 (0.316 ≤ |*ρ*| ≤ 0.803, p < 0.05). The correlation with the MN recruitment threshold exactly mirrors the correlation with the discharge frequency.

#### Population based analysis

We performed a proof-of-concept investigation into whether the distribution of reflex amplitudes can be used to infer the distribution of MN properties. Therefore, we estimated reflex amplitudes of the simulated MN pool for five different distributions of MN size parameters and two activation levels. Figure 5 shows the empirical cumulative probability for MN input resistance and reflex amplitude estimated from PSF and PSTH. MN pools with proportionally more small MNs show a bias towards higher reflex amplitude values in the PSF (Figure 5, blue). Accordingly, MN pools with proportionally more large MNs show a bias towards lower reflex amplitudes in the PSF (Figure 5, red). This pattern can be observed for both activation levels 3 and 5. In contrast, the cumulative probability distributions of PSTH reflex amplitudes mostly overlap for all MN pools at activation level 3. At activation level 5, the MN pool with proportionally more small MNs shows the lowest reflex amplitudes in PSTH (Figure 5B, blue). MN pools with proportionally more large MNs cannot be distinguished based on their PSTH reflex amplitudes (Figure 5B, yellow, orange and red).

**Figure 5.**
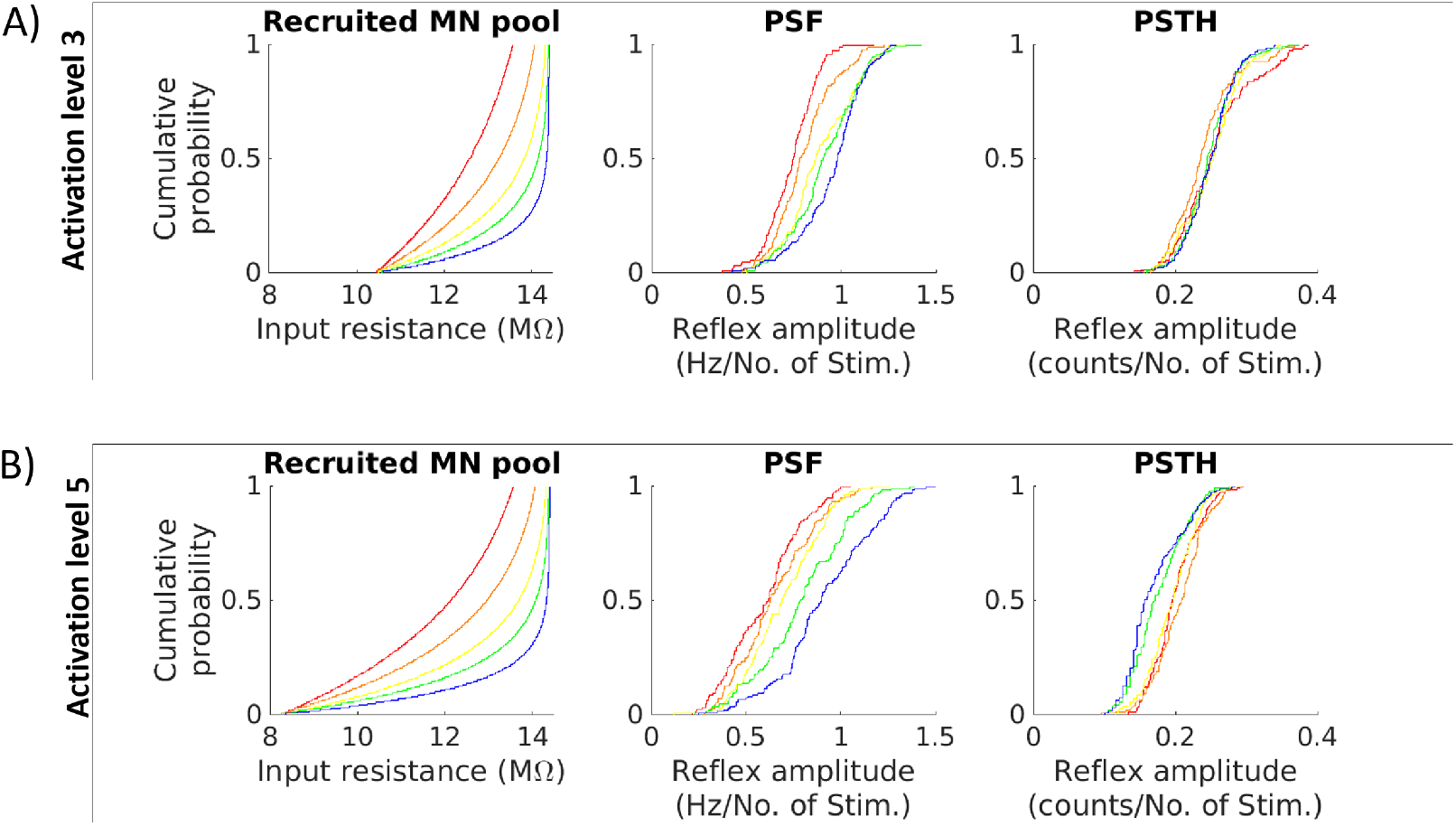
Empirical cumulative probability of MN pool reflex amplitudes for different simulation settings. Shown are results obtained with different MN pool compositions, represented by the input resistance. The cumulative probabilities of the respective reflex amplitude values estimated from PSF and PSTH are shown in similar colors. In (A), activation of level 3 was applied, while the results in (B) were obtained with activation level 5. Yellow corresponds to the MN pool used in the above analyses.

## 3 Discussion

The MU reflex profile is commonly used to estimate the EPSP size and reflex excitability of MUs in association with their phenotype (Binboğa and Türker, 2012; Awiszus and Feistner, 1993). This view is mainly based on the assumption of an MN as a linear integrator. However, in the present study, we found that the reflex amplitude variation among a (heterogeneous) population of MUs is affected nonlinearly by several factors, e. g., DR, membrane noise, and MN properties. This observation aligns with previous theoretical and experimental findings, i. e., that DR and background noise affect the firing probability of individual MUs (Jones and Bawa, 1997; Matthews, 1999; Herrmann and Gerstner, 2002; Piotrkiewicz et al., 2009; Türker and Powers, 1999). When fixing the stimulus intensity, the variability in the reflex amplitude can not be explained by the phenotype and discharge regime of the MU only. Due to this multivariable uncertainty in *in-vivo* experimental data, examining factors contributing to the variation in reflex amplitude is not straightforward and requires a modelling approach.

### Experimental findings

We reconstructed a large population of MU spike trains (N=206) during H-reflex measurements by decomposing both high-density surface EMG and intramuscular EMG signals. The method was previously validated for similar isometric contraction forces (Yavuz et al., 2015).

The *in-vivo* results demonstrated a dependency of reflex amplitude in several individual MUs on their DR and CoV_ISI_. The linear regression between DR and reflex amplitude was predominantly positive at 10 % MVC when the PSF method was used to measure reflex amplitudes. This result suggests that the background DR of a MN is one of the determinants of PSF trajectory across the population, just like shown in single MNs (Türker and Powers, 1999). As shown earlier, the PSF can provide information about the PSP amplitude given that the firing rate is linearly related to the amplitude of the membrane potential at the MN soma (Türker, 2022). However, this linear relation can only be assumed within a limited scope, e. g., for MNs with very similar properties and at the same force. The relationship between DR and reflex amplitude became inconsistent as the contraction force increased to 20 % MVC. This variability is likely attributable to changes in overall membrane noise with increasing contraction force (Kudina, 1999; Moritz et al., 2005), which, in turn, affects the probability of reflex excitation (Türker and Powers, 1999). The PSTH method, on the other hand, did not show any significant dependency on either DR or ISI variability. The discrepancy between the two methods will be discussed further in the following section.

While these results showed the extent to which discharge statistics account for the reflex amplitude variation, it is important to acknowledge the direct role of the unknown afferent input distribution across the MN pool (Binder et al., 2002). Previous research paints a heterogeneous picture of the distribution of Ia monosynaptic afferent inputs to the MN pool. For example, Binboğa and Türker (2012) found that the H-reflex amplitude is higher in larger MUs of the human soleus muscle, while Awiszus and Feistner (1993) found the opposite with a similar experimental setup.

### Simulations reveal multivariable influence on reflex amplitudes

We investigated the variation in the reflex amplitude of a population of MNs using a computational model where the influencing factors were reduced to the intrinsic properties of MNs and the composition of synaptic input. We simulated a pool of 200 MNs with membrane resistances distributed exponentially, resulting in a population containing many low-threshold and relatively fewer high-threshold MNs. The total input current was limited to a uniformly distributed common and independent noise. As discussed before, the distribution of Ia afferent input is a critical but unknown parameter that can directly determine the reflex distribution. Consequently, many modelling studies adopt both the uniform (e. g., Elias et al., 2014; Lin and Crago, 2002) and biased Ia input distribution, where larger MUs have higher input gain (e. g., Dideriksen et al., 2015). Regardless of these perspectives, we used a uniform Ia input distribution in this study to control for synaptic input variation and investigate the effect of discharge statistics on reflexes across the population.

In the PSF, the number of MNs with significant reflex amplitude decreased with the increased activation level compared to the total number of recruited MNs. The amplitude of the noise input was scaled proportionally to the mean input while the reflex stimulus remained constant. We conclude that the reflex response becomes less distinguishable from noise with increasing background activity.

We found a positive correlation between DR and reflex amplitude measured using PSF in most activation levels. Only the regression at the lowest activation level was negative, most likely due to the number of MNs close to their recruitment thresholds.

The PSF trajectory of these MNs is not correlated with the EPSP trajectory because of their non-stationary firing regime (Türker and Powers, 2005). Similar to the experimental results, the positive linear regression between DR and the reflex amplitude was diminished when we used the PSTH method to measure the reflex amplitude. However, when eliminating the membrane noise, the PSTH showed a size-related pattern as well. This finding suggests that the interpretation of reflex amplitude variation may change based on the mathematical foundation of the analysis method and the experimental conditions, as discussed below.

### Disparity between the PSF and PSTH results

The PSTH and PSF are two well-established methods for investigating single MU reflex trajectories. However, a debate spanning several decades concerns whether the reflex trajectory measured by either method can reliably estimate the EPSP. We compared the distribution of reflex amplitudes determined using two metrics, PSTH and PSF, across experimentally sampled and simulated MNs. We found a disparity between the PSTH and PSF results in their interpretation of reflex amplitude variation across the MN pool. The simulation results provided additional insight into the understanding of this disparity.

Analyzing the PSTH measurements, we found no dependency between reflex amplitude variability and MN size. Namely, the reflex amplitude among MNs decreased evenly with increasing activation level across all MNs until it reached a plateau. This pattern aligns with previous modelling studies using PSTH (Jones and Bawa, 1997). Matthews (1999) and Herrmann and Gerstner (2002) reported similar trajectories. The PSTH is less responsive to variation in DR since the PSTH is a probability-based estimation and probability during the reflex does not change rigorously when the EPSP timing changes slightly with respect to the last action potential (Türker and Powers, 1999). Since the PSF reflects absolute changes in the DR, the timing of the reflex stimulus with respect to the current phase of the membrane potential trajectory will be more prominently reflected in the PSF (Türker, 2022). Indeed, the PSF results demonstrated a significant correlation between reflex amplitude and MN sizes. This correlation was stronger at higher input currents (force) levels. This trajectory shows similarity with the slope of the current-frequency relation, especially for the smaller MNs. The slope of their current-frequency relation rises more steeply just after recruitment, takes a smaller and approximately constant slope for a large range of input currents and then increases again for even larger inputs (Figure 3A).

Based on these results, it can be concluded that the PSF method is susceptible to heterogeneity across the MN pool and the nonlinear combination of the DR and membrane noise of the MN that dictates the size of the excitatory reflex. Therefore, not only the type of MN and the corresponding distribution of the Ia input but also the nonlinear integration process of those MNs should be taken into account to explain the variation in the reflex amplitude based on the PSF.

The variation in the measured reflex amplitude can be advantageous when investigating heterogeneity within a population, as it amplifies the disparity between different types of MUs, even though the variation is nonlinear. Indeed, the simulation results demonstrated a strong correlation between MN size and reflex size as measured by PSF. The present experimental study further found that higher ISI variability, used as a measure of total membrane noise, may act as an additive factor introducing nonlinearity into the system by modulating the influence of DR on reflex amplitude variation. Indeed, the simulation results confirmed that the ISI variability is a prevalent factor in the PSTH that wipes out the effect of MN size. In the PSF, on the contrary, we observed a shift in the variability with respect to the activation level. We attribute this behaviour to a decrease in the overall excitatory input as a result of missing membrane noise. A similar finding has been reported for single MU measurements (Türker and Powers, 1999).

### Population statistics

Taken together, these results highlight the need to choose appropriate metrics when using MU reflex discharges as indicators of reflex excitability. The multifactorial variation in reflex discharge complicates using the reflex amplitude as a predictor of reflex excitability across a population of spinal MNs. The comparison of the estimated reflex amplitude across different experimental conditions may show significant differences unrelated to the excitability and amplitude of excitatory or inhibitory synaptic inputs to the MNs. However, the uncertainty due to the variation in reflex amplitude can be reduced by considering the proportional behaviour of a sampled population instead of the absolute reflex amplitude of individual MUs. For this, PSF is a suitable method as it better reflects the heterogeneity within the sampled MUs.

The simulation results showed that the shape parameters of the probability density of reflex amplitude well reflect the change in the distribution of MN size when keeping other influencing parameters constant. This result is in line with our previous findings, where we showed that the shape of the probability density function changes with the gain of the reflex transmission measured *in vivo* (Yavuz et al., 2018, 2015). These are preliminary results that point the way for further research. Variability can be investigated more thoroughly using e. g., sensitivity analysis or bootstrapping methods. Extracting a suitable metric from these results is beyond the scope of this work. However, we speculate that estimators of the shape parameters (e. g., skewness, kurtosis) can be crucial metrics to tackle the variability problem since they are non-dimensional distribution quantities. They should be less affected by the variation of reflex amplitude due to varying input current, stimulation intensity, or firing statistics than scale parameters (Ayachi et al., 2014). Therefore, a method based on the shape of the probability density function can potentially provide a reliable solution to investigate spinal adaptation and its neural mechanisms, an unsolved challenge for several decades.

### 3.1 Limitations

In this study, MUs were sampled *in vivo* at 10 % and 20 % MVC. Consequently, we assumed that population heterogeneity was primarily limited to low-threshold MUs. The contraction force was capped at 20 % MVC for two main reasons: first, to ensure high decomposition accuracy, and second, to minimize fatigue as an additional factor influencing reflex amplitude variability. For the same reason, the number of stimulations was kept between 100 and 150. However, our preliminary results indicate that the error in both PSF and PSTH cusum decreases exponentially with the number of stimulations and reaches a plateau only after approximately 200 stimulations (Schmid, 2024).

We chose a computational model that provides a good estimate of motor neuron responses to changes in effective synaptic inputs (Negro and Farina, 2011). Limiting the space of influencing factors, we excluded any modulatory current, like PIC. We collapsed the dendrite into a single compartment, thereby considering the contribution of the dendrite to the overall input resistance. Due to their slow time dynamics, we expect no major changes in PIC activity during short stimuli (Binder et al., 2020). However, since PICs modulate the background DR, they do have an indirect effect on the estimated reflex amplitude. The MN model does not consider threshold variations within an ISI. The spike threshold was shown to vary within the ISI in a way that it follows the membrane potential trajectory of the AHP (Calvin, 1974; Powers and Binder, 1996). In the employed model, the threshold plateaus towards the end of the ISI. Thus, we might overestimate the efficacy of stimuli delivered late in the interspike interval. The effect of the spike threshold on reflex amplitudes could be investigated by adapting a spike-response model, as e. g., proposed by Herrmann and Gerstner (2002).

Finally, note that we only used excitatory stimuli. A recent work highlights subject-specific variability in inhibitory spinal circuits (Pascual-Valdunciel et al., 2025). Even though our results are not directly transferable to inhibitory reflexes, we additionally expect intra-subject variability due to the heterogeneity of the MU population for inhibitory reflexes. Considering previous studies, which suggested differences between PSTH and PSF for inhibitory stimuli (Türker and Powers, 2005, 1999), further investigations on inhibitory reflexes are required.

### 3.2 Conclusion

Analyzing reflex amplitudes at the population level can be essential for understanding how reflex excitability is distributed within a heterogeneous MU population. This approach is particularly important for identifying excitability patterns that may underlie functional adaptations or pathological changes in the neuromuscular system. This study provides valuable insights that highlight the importance of considering variability when evaluating and interpreting *in vivo* MU reflex data. The most important implications are: (1) The PSTH is sensitive to membrane noise but more robust against experimental conditions and is well suited for investigating the reflex responses of the MN pool as an entity and for comparing different experimental conditions, like force levels. (2) The PSF is sensitive to the MN size and is well suited for investigating differences in reflex amplitudes within a MN pool, but can be affected by inter-subject differences. (3) The uncertainty due to the variation in reflex amplitude can be reduced by using the probability of a large sample from the entire population. Together with methods that enable recording from a large number of MUs, population-based analyses can broaden our understanding of MUs and spinal circuits.

## 4 Methods

### 4.1 *In-vivo* study

To investigate the above-mentioned hypothesis, we utilized *in-vivo* MU data previously published by Yavuz et al. (2015). The experimental setup and procedure described in Yavuz et al. (2015) are outlined below.

#### Ethical statement

The protocols were approved by the Human Ethics Committee of the University Medical Center, Georg-August-University of Göttingen (approval date: 1/10/12). Subjects provided written informed consent before the experiment.

#### Experimental setup

Data were obtained from six healthy subjects (males, age: 29.6 ± 5.6 years). Subjects were seated on the chair of a dynamometer system (Biodex Medical Systems Inc., NY, USA) with their right foot and leg fastened. The ankle angle was arranged to the anatomical rest position of the subject (approx. 10^◦^ plantar flexion), and the subject chair was arranged such that the knee angle was 120^◦^ (Yavuz et al., 2014). Isometric dorsiflexion torque was measured through a transducer (ATI Omega160) mounted to the footplate of the dynamometer. High-density surface EMG (HDsEMG) and intramuscular wire electrodes were used to identify the spiking activity of MUs. By combining surface and intramuscular EMG, we aimed to sample MUs from different muscle depths. First, two bipolar Teflon-insulated silver wire electrodes (75 µm in core diameter) were inserted into different locations of the tibialis anterior (TA) muscle. The HDsEMG electrodes (5×13 electrode grid, 8 mm interelectrode distance, ELSCH064NM2, OT Bioelettronica, Torino, Italy) were placed parallel to the tibia and covered the belly of the TA (Yavuz et al., 2015). An ankle strap electrode was used as a reference electrode. The common peroneal nerve (CPN) was stimulated with the anode of the stimulation probe (a dampened 12 x 8 cm pad, Spes Medica s.r.l., Battipaglia, Italy) posterior of the fibula’s head. The metal cathode of the stimulation probe was placed anterior to the fibula’s head (Figure 6A). Signals were recorded at 10240 sample/s using an EMG-USB2 data acquisition system (OT Bioelettronica, Torino, Italy). The intramuscular and the HDsEMG signals were filtered by a built-in band-pass filter with 100–4400 Hz and 10–500 Hz cut-off frequencies, respectively.

**Figure 6.**
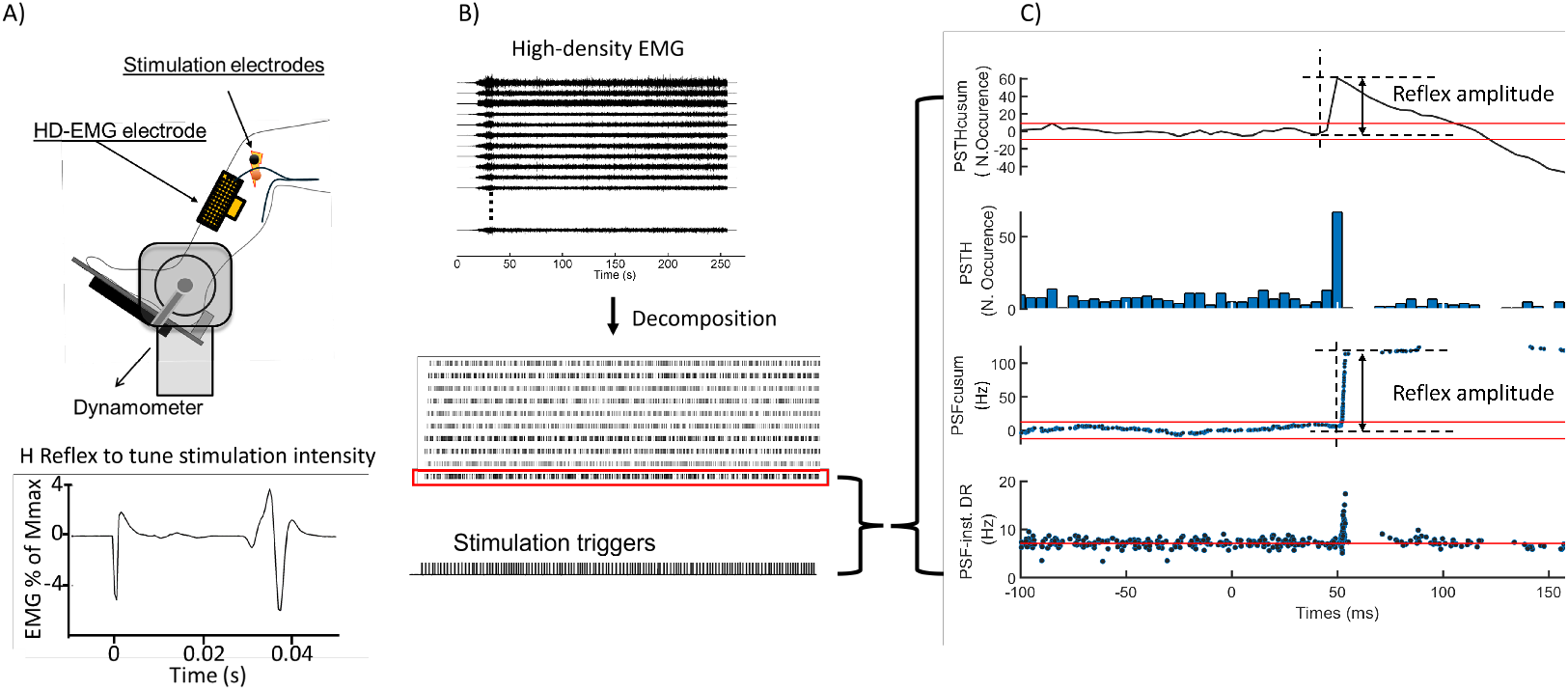
Illustration of the experimental setup and protocol to tune stimulation intensity (A), the decomposition process (B), and the reflex amplitude measurement from PSF- and PSTH-cusum graphs (C).

#### Procedures

Subjects performed a familiarisation contraction before the protocol started. After tuning the stimulation intensity to elicit an H-reflex with a size equal to 10 % of the maximum M-wave (Figure 6A), subjects were asked to perform sustained isometric ankle dorsiflexion contractions at 10 % and 20 % of the maximum voluntary contraction force (MVC). During the sustained contractions, about 200 electrical stimuli (100 µs pulse duration) with 2-3 s random inter-pulse intervals were delivered to the CPN.

### 4.2 Simulation study

#### Motor neuron pool model

We established a computational MN pool model to investigate the H-reflex variation in controlled conditions. The model contained 200 MNs, comparable to the number of MNs innervating the TA muscle (Duchateau and Enoka, 2022). We prescribed the membrane resistance and input current distributions to be able to distinguish the effect of discharge properties on reflex amplitude variation among a large population of MNs. The distribution of the membrane resistance was exponential. The distribution of monosynaptic spindle afferent input to MNs was uniform to eliminate variation due to input current. MNs were simulated using a two-compartment model previously proposed (Negro and Farina, 2011). The membrane potential in the soma compartment consists of three voltage-dependent conductances (mediating Na^+^-, fast K^+^-, and slow K^+^-currents) described by four gating variables, which are functions of the voltage-dependent rates (Cisi and Kohn, 2008; Negro and Farina, 2011). For details on the model, we refer to the published code (https://github.com/IMSB-CBM/ReflexAmplitudeEstimation.git).

The MN input comprised a common and an independent component (Figure 7). The common component was represented by a constant mean value and a zero-mean band-pass filtered (fourth-order butterworth filter) Gaussian noise (15 Hz to 35 Hz, Conway et al., 1995; Halliday et al., 1998; Negro and Farina, 2011), mainly determining the discharge frequency and similarly applied to all MNs. The constant mean input was varied from 4 nA to 16 nA to simulate different activation levels, which correspond to the scale of muscle force used during the experiments. The independent input component was individually modeled for each MN as zero-mean, low-pass filtered (< 100 Hz, second-order butterworth filter) Gaussian noise (Negro and Farina, 2011). We fixed the standard deviation of the common noise to 20 % of the mean input, and the standard deviation of the independent input was scaled to account for 20 % of the total noise (Negro et al., 2016b). This choice yielded values of 10 % < CoV_ISI_ < 30%, which are within the physiological range (Matthews, 1996; Moritz et al., 2005).

**Figure 7.**
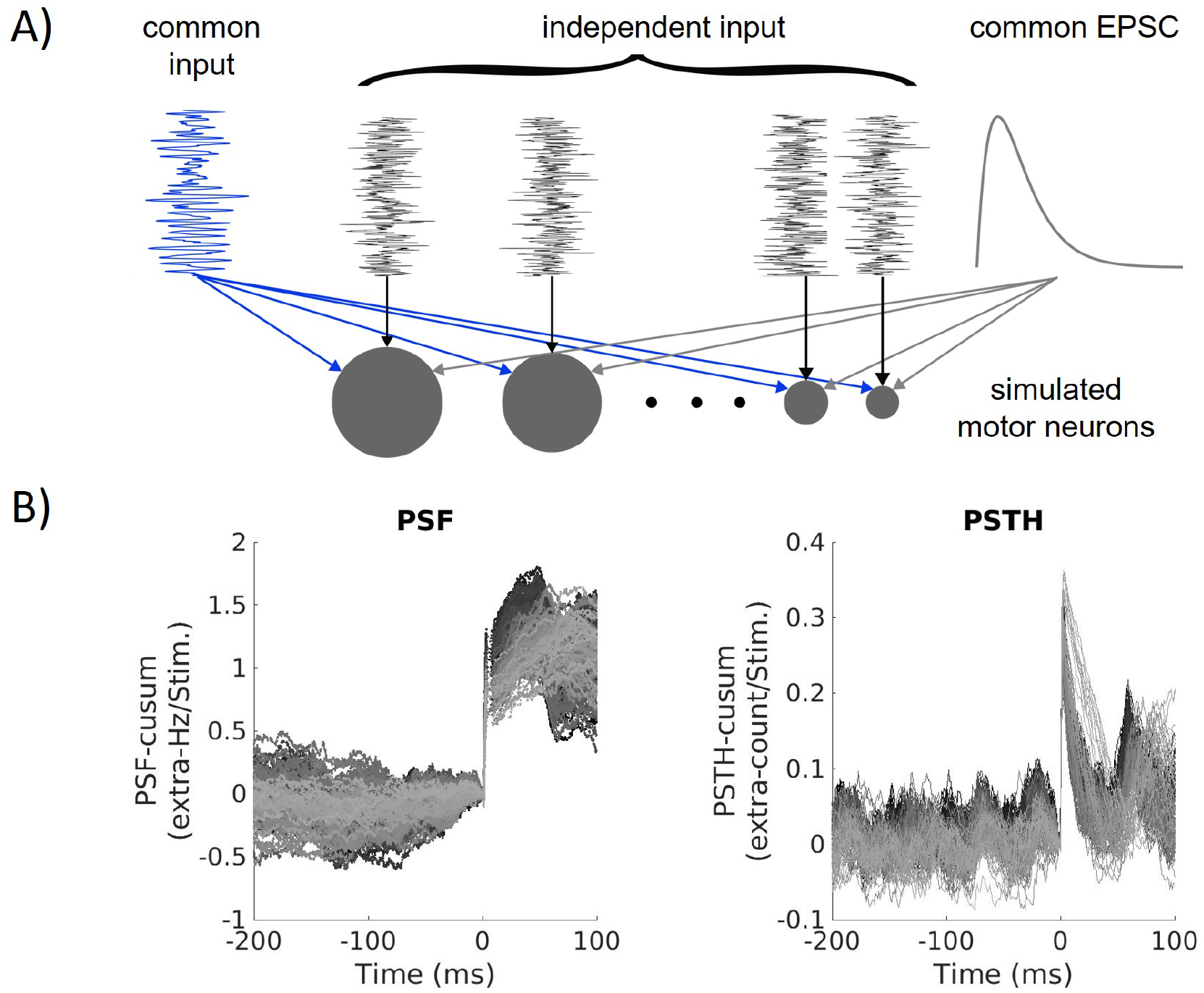
A: Schematic drawing of the inputs to the simulated motor neuron (MN) pool. Each MN (grey circle) receives a common input (blue) and an excitatory postsynaptic current (EPSC, grey), which are similar for all MNs. In addition, each MN receives an individual input (independent noise, black). Note that in the figure, the inputs are not scaled with respect to each other. B: PSF- and PSTH-cusum of the simulated MN pool (shown is a 300 ms excerpt of the data for better visibility). Each line represents one MN.

We modeled the reflex stimulus as an excitatory postsynaptic current (EPSC)

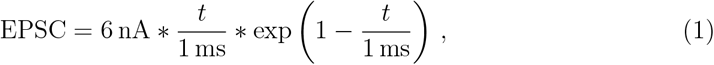

and assumed a stimulus duration of 40 ms. The time constant of 1 ms was chosen to match EPSC rise times measured in previous studies (Lüscher et al., 1983; Hochman and McCrea, 1994). The gain of EPSC (6 nA) was chosen empirically to obtain reflex sizes comparable to the experimental data. By applying a fixed current, we account for the nonlinear properties of the membrane, resulting in different-sized EPSPs depending on the current value of the membrane potential.

In each simulation, the stimulus was applied 200 times with normally distributed interstimulus intervals of 1000 ± 100 ms. All inputs were linearly summed and applied to the soma compartment of the MN model, i. e., representing effective synaptic currents (Heckman and Binder, 1988). In the simulation, we refrain from providing MVC estimates to avoid making assumptions about the MU twitch properties. Instead, we use unitless activation levels that consider the number of recruited MNs. We divided the effective synaptic input into seven different segments each of which corresponds to an arbitrary activation level. Depending on the muscle, full recruitment is achieved between 50 % and 80 % MVC (Moritz et al., 2005; De Luca et al., 1982; Kukulka and Clamann, 1981). Based on that we assumed that the activation levels 2 to 4 correspond to 10 and 20 % MVC, which we used in experimental research. All simulations were performed with MATLAB R2021a (9.10.0.1602886).

### 4.3 Data analysis

#### Experimental data

Single MUs were identified in HDsEMG signals using a convolutive blind source separation method (Negro et al., 2016a). The accuracy of HDsEMG decomposition in the case of electrical stimulation was extensively validated in a previous study (Yavuz et al., 2015). The MU spike trains with silhouette score lower than 80% (SIL < 80 %), CoV_ISI_ higher than 30 %, and background DR lower than 5 Hz were eliminated (Yavuz et al., 2018). The intramuscular EMG (iEMG) signals were decomposed into MU action potentials using EMGLAB (McGill et al., 2005). The decomposition results were visually inspected and manually edited through the editing interface of EMGLAB. Later, we determined MUs commonly identified from iEMG and HDsEMG, estimating the number of matched discharges with the discharge timing tolerance set to ±5 ms (Yavuz et al., 2015). Either of the common MUs was eliminated to avoid duplicated spike trains. The reflex responses of simulated and *in-vivo* MUs were examined through PSF and PSTH methods. Both PSF and PSTH were computed for 300 ms time windows around stimulation instant (200 ms before and 100 ms after stimulation). Nevertheless, determining reflex activity in raw PSF/PSTH is difficult and may not be accurate enough due to instantaneous discharge variability. Therefore, we measured reflex parameters (latency, duration, and amplitude) from the cumulative sum (cusum) of PSF and PSTH (Figure 6B, C). The reflex amplitude was measured as the difference between the onset of the reflex response and the first turning point before the peak values in PSF-cusum and PSTH-cusum (Figure 6C). Only values exceeding the error interval (100 % of the maximum pre-stimulus cusum deflection from zero) were considered as a significant reflex response (Türker et al., 1997). The amplitude value was normalized by the number of stimulations (Miles et al., 1989). Through normalization, we described the extra-discharge or extra-count per stimulation. The mean DR of each MU was calculated from the pre-stimulus time interval of 500 ms. The relationship between the simulated MU size (represented by input resistance) and the reflex amplitude was investigated by comparing the shape characteristics of the cumulative probability distributions of reflex amplitude and input resistance values.

#### Automated evaluation of reflex amplitudes for simulations

As with the experimentally recorded MUs, reflex amplitudes of simulated MNs were also obtained from PSF- and PSTH-cusum. Spike trains of the simulated MNs were directly obtained from the simulated membrane potential trajectory. Simulated MNs with background DR lower than 7 Hz and CoV_ISI_ higher than 30 % were not considered for the analysis (considering that the model is fitted to cat data, that is a comparable range).

In this work, reflex amplitudes of several hundred simulated MNs under numerous conditions were determined. For this large amount of data, manual evaluation is not feasible, and an automated reflex-amplitude estimation algorithm was employed. Similar to the manual evaluation, an error box approach was chosen to determine significant reflex responses. Only responses exceeding the largest pre-stimulus deflection of the cusum from zero were considered significant reflex responses. For the simulation data, we used a pre-stimulus time window of 300 ms and a post-stimulus time window of 150 ms. The onset and end of the reflex response were determined using the “steepness” of the reflex response, determined by the forward difference of the cusum. Therefore, a threshold was determined, similar to the cusum error box. The largest absolute deflection from zero of the approximated slope of the cusum defines the reflex threshold, and any point above the threshold is considered to be part of the reflex response.

Further, we only considered reflex responses up to 15 ms after stimulus time. Since we did not consider conduction delays, the reflex response must occur in this time period. To further prevent faulty assignments of reflex amplitudes, the results of the automated evaluation were also visually inspected.

#### Statistical analysis

The reflex parameters (latency, duration, and amplitude) and MU discharge features (mean DR and CoV_ISI_) across different contraction forces were analyzed using one-way ANOVA. Linear regression analysis was performed to assess the effect of DR and CoV_ISI_ on reflex variability. The relationship between DR and reflex amplitude was evaluated for each subject and contraction force using both the PSF and PSTH methods. Additionally, the regression between CoV_ISI_ and reflex amplitude was estimated by pooling MUs, selected based on the significance of the DR–amplitude regression model. Group 1 comprised subjects 2, 3, and 4, who exhibited a significant positive regression at 10 % MVC, while Group 2 included subjects 1, 5, and 6, who showed no significant regression. To compare the relative importance of each coefficient in a regression model, we standardized the amplitude values estimated z-score 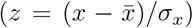. The correlation between the simulated MN’s reflex amplitudes and threshold and DR, was estimated using a bivariate Pearson correlation test (one tail). In all the tests, the significance level of *p* < 0.05 was chosen.

## Author Contributions

**Conceptualization:** Laura Schmid, Thomas Klotz, Francesco Negro, Utku Ş. Yavuz.

**Investigation:** Laura Schmid, Utku Ş. Yavuz, Francesco Negro.

**Methodology:** Laura Schmid, Utku Ş. Yavuz.

**Software:** Laura Schmid.

**Visualization:** Laura Schmid.

**Formal analysis:** Laura Schmid, Thomas Klotz, Oliver Röhrle, Christopher K. Thompson, Francesco Negro, Utku Ş. Yavuz.

**Writing - original draft:** Laura Schmid, Thomas Klotz, Utku Ş. Yavuz.

**Writing - review & editing:** Laura Schmid, Thomas Klotz, Oliver Röhrle, Christopher

K. Thompson, Francesco Negro, Utku Ş. Yavuz.

## Acknowledgements

This research is supported by the German Research Foundation (Deutsche Forschungs-gemeinschaft, DFG) as part of the priority program SPP 2311 (465243391, to OR), the European Research Council (ERC) through ERC-AdG “qMOTION” (101055186, to OR), and Consolidator Grant INcEPTION (101045605, to FN), and the National Institutes of Health (NIH) through grants NS125863 and NS124820 (to CKT).

## Data availability statement

The scripts to run the simulations are published in a repository under https://github.com/IMSB-CBM/ReflexAmplitudeEstimation.git.

